# Telomere Dysfunction Drives Chronic Lung Allograft Dysfunction Pathology

**DOI:** 10.1101/746768

**Authors:** Ram P. Naikawadi, Gary Green, Kirk D. Jones, Natalia Achtar-Zadeh, Julia E. Mieleszko, Jasleen Kukreja, John Greenland, Paul J. Wolters

**Affiliations:** Division of Pulmonary, Critical Care, Allergy and Sleep Medicine, Department of Medicine, University of California, San Francisco, CA.; Department of Pathology, University of California, San Francisco, CA.; Department of Surgery, University of California, San Francisco, CA.; Medical Service, VA Health Care System, San Francisco, CA

**Keywords:** Airway fibrosis, Telomere, Club cells

## Abstract

**Rationale:** Telomere dysfunction is associated with multiple fibrotic lung processes, including chronic lung allograft dysfunction (CLAD) which is a major limitation to long-term survival following lung transplantation. Although shorter donor telomere lengths are associated with an increased risk of CLAD, it is unknown whether short telomeres are a cause or consequence of CLAD pathology.

**Objective:** Our objective was to test whether telomere dysfunction contributes to pathologic changes seen in CLAD.

**Methods and Results:** Histopathologic and molecular analysis of human CLAD lungs demonstrated shortened telomeres in lung epithelial cells quantified by teloFISH, increased numbers of surfactant protein C immunoreactive type II alveolar epithelial cells (AECs), and increased expression of senescence markers (beta-galactosidase, p16, p53 and p21) in lung epithelial cells. Telomere repeat binding factor 1 flox/flox (*TRF1^F/F^*) mice were crossed with tamoxifen inducible SCGB1a1-cre mice to generate *SCGB1a1-creTRF1 ^F/F^* mice. Following 9 months of tamoxifen-induced deletion of TRF1 in club cells, mice developed mixed obstructive and restrictive lung physiology, small airway obliteration on micro-computed tomography, a 4-fold decrease in telomere length in airway epithelial cells, collagen deposition around bronchioles and adjacent lung parenchyma, increased type II AEC numbers, expression of senescence-associated beta-galactosidase in epithelial cells and decreased SCGB1a1 expression in airway epithelial cells.

**Conclusions:** These findings demonstrate that telomere dysfunction isolated to club cells leads to airway-centric lung remodeling and fibrosis similar to that observed in patients with CLAD and suggest that lung epithelial cell telomere dysfunction may be a molecular driver of CLAD.

## INTRODUCTION

Lung transplantation is a lifesaving therapy for patients with end-stage lung disease. Chronic lung allograft dysfunction (CLAD) is a common complication that limits long-term survival after lung transplantation (1) with 50% of patients succumbing to CLAD within 5 years (2). CLAD is a syndrome clinically defined primarily by a decline in lung function. CLAD is subdivided into obstructive and restrictive subtypes, known as bronchiolitis obliterans syndrome (BOS) and restrictive allograft syndrome (RAS), respectively (3). While these subtypes may overlap, they have characteristic pathologies and imaging findings. BOS is characterized pathologically by constrictive or obliterative bronchiolitis, while RAS is characterized by a variable combination of constrictive bronchiolitis, pulmonary fibrosis, and pleuroparenchymal fibroelastosis (4, 5).

While immune responses are thought to play a role in CLAD pathogenesis, augmentation of immunosuppression is generally not effective in reversing CLAD progression (6). Thus, understanding molecular mechanisms underlying the pathologies of CLAD may be useful for understanding this syndrome. Airway epithelium consists of club cells, ciliated cells, basal cells, and neuroendocrine cells. Of these, club cells show particular evidence of damage in patients with CLAD (7). In addition to secreting protective factors, club cells replicate and differentiate to replenish airway epithelial cells and thus may be under stress in the context of lung transplantation. As a potential marker of replicative stress (8), short telomeres in lung airway epithelial cells have been associated with development of CLAD (9). Further, explanted lung tissue from patients with CLAD show evidence of telomere shortening (10).

Telomere dysfunction manifests clinically as fibroproliferative lung pathologies of which usual interstitial pneumonia is the most common, but other patterns of fibrosis are also observed (11). Conditional deletion of telomere shelterin proteins in specific cell types has been shown to model pathologies of human diseases mediated by telomere dysfunction (12–15). For example, modeling telomere dysfunction in surfactant protein C-expressing cells results in lung fibrosis. To test the hypothesis that telomere shortening in club cells is sufficient to induce CLAD pathology, we developed a mouse model where the shelterin protein family member Telomere Repeat Binding Factor 1 (TRF1) was selectively deleted in club cells.

## METHODS

### Human specimens

Fibrotic lung tissue was obtained at the time of lung transplantation from patients with a pathological diagnosis of chronic lung allograft dysfunction. Control human lungs consisted of lungs not used by the Northern California Transplant Donor Network; our studies indicate that these lungs are physiologically and pathologically normal (16). Endobronchial and transbronchial biopsies were performed within 90 days post-transplant and sections were recut from FFPE tissue blocks, as previously described (9). Written informed consent was obtained from all subjects and was approved by the University of California, San Francisco Institutional Review Board protocol number 13-10738.

### Mice

*TRF1^F/F^* mice were purchased from Jackson lab, *SCGB1a1-cre/ERT* mice were provided by Jason Rock, at UCSF. Mice were bred and housed in pathogen-free conditions in accordance with the guidelines of Laboratory Animal Resource Center (LARC). All animal procedures were carried out using protocols approved by the Institutional Animal Care and Use Committee (IACUC) at the University of California, San Francisco.

### Tamoxifen administration

Tamoxifen (Toronto Research Chemicals T0600), suspended in peanut oil was injected (250 mg/kg body weight once per week) intraperitoneally to *SCGB-creTRF1^F/F^* mice and *TRF1^F/F^* littermate controls beginning at 10 weeks age.

### Hydroxyproline assay

Hydroxyproline levels were quantified as described previously (17).

### Telomere Q-FISH assay

Telomere lengths were measured on paraffin-embedded tissue sections by Quantitative Fluorescence in Situ Hybridization (Q-FISH) as described (15).

### Senescence Associated-β-galactosidase (SA-β-gal) assay

To test for SA-β-gal (18) staining on tissues following protocol was used as described (19).

### Lung Mechanics

Lung mechanics were measured as described by the manufacturer (FlexiVent; SCIREQ Inc, Canada). To measure pulmonary physiology mice were anesthetized with i.p. ketamine/xylazine and adequate depth of anesthesia was assured by observing pedal withdrawal reflex. A tracheostomy was performed on the mouse in deep-state anesthesia. A tubing adaptor (20 gauge) was used to cannulate the trachea. The mouse was then attached to a rodent ventilator and pulmonary mechanics analyzer and ventilated at a tidal volume of 9 ml/kg, a frequency of 150 breaths/minute, and 3 cm H2O positive end-expiratory pressure. The mouse was paralyzed with pancuronium bromide (0.1 mg/kg intraperitoneally) to facilitate static compliance measurement. before tracheal cannulation and pancuronium administration. Lung mechanics were measured using forced oscillation maneuver algorithms using flexivent software.

### Statistics

Statistical analysis were performed with GraphPad Prism software (Version 7.0) P values were calculated using two-tailed Student’s *t* test. Welch’s correction was applied for analysis of unpaired groups. Differences in survival were analyzed by *log-rank (Mantel-Cox)* test. Data are represented as the mean ± SEM. *P* values represent the significance using the following symbols. ns = not significant, p>0.05, *p<0.05, **p<0.01, ***p<0.001, ****p<0.0001.

## RESULTS

### Histopathology of CLAD lungs

Masson’s trichrome staining on sections of explanted lung from human lung transplant recipients with CLAD revealed increased collagen deposition around bronchioles (Fig. 1A), parenchymal fibrosis as quantified by the Ashcroft scoring method (20) (Fig. 1D), and concentric narrowing of airway lumens compared to normal lungs. Patchy bronchiolar obliteration was also observed in CLAD lungs (Fig. 1A). Moreover, immunofluorescence staining identified alpha smooth muscle actin-expressing mesenchymal cells around bronchioles in CLAD lungs, unlike normal lungs (Fig. 1B). Immunostaining for surfactant protein C (SPC) revealed type II alveolar epithelial cell (AEC) hyperplasia (Fig. 1C) with up to 5-fold more type II AECs (Fig. 1E) adjacent to bronchioles.

**Fig. 1.**
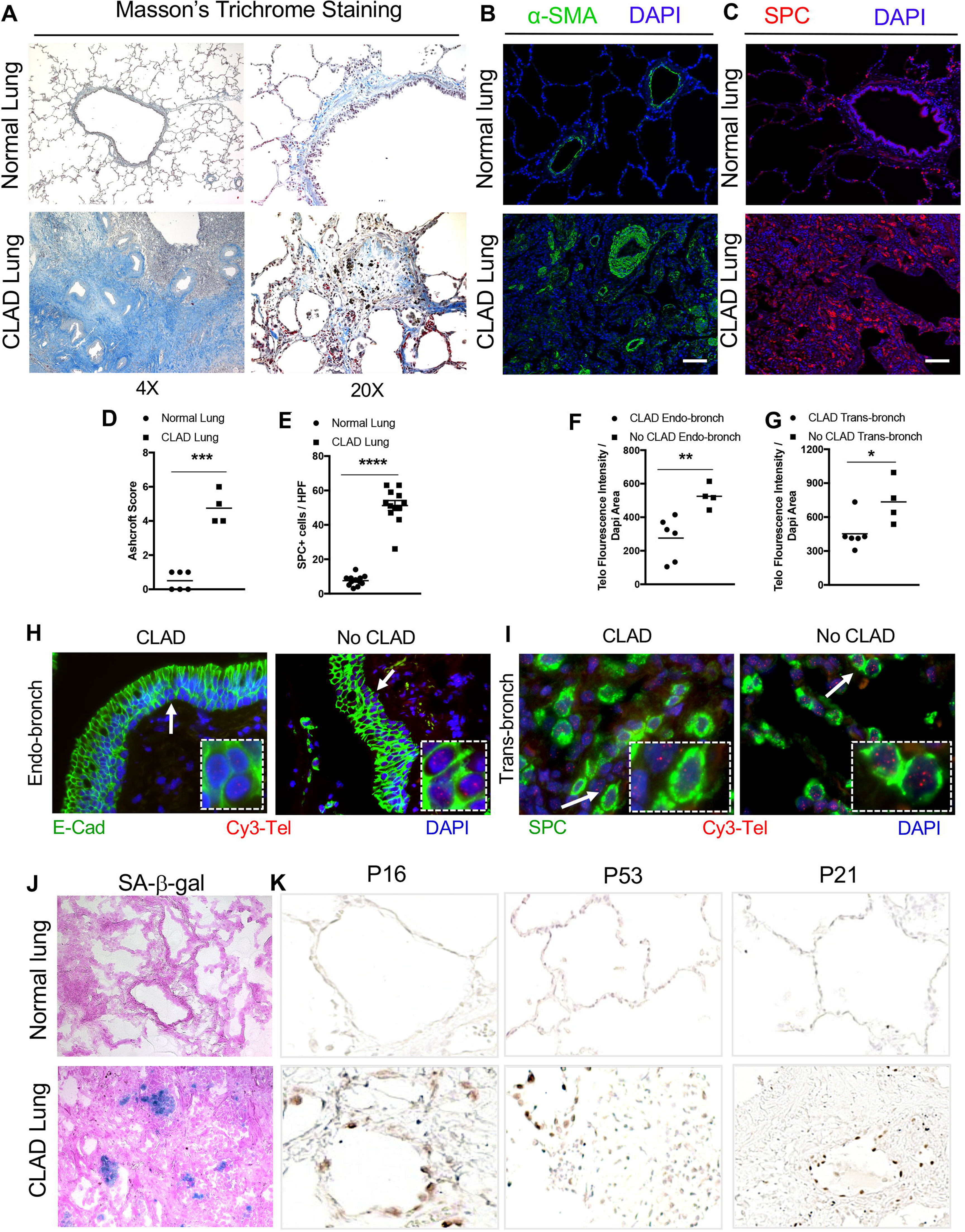
CLAD pathology in lung transplant recipients. **(A)** Masson’s trichrome staining in CLAD lungs and Normal lungs. Collagen deposition (blue) is seen around the airways and in bronchiole causing obliteration in CLAD lungs. Magnification 4X (left panels) and 20X (right panels) **(B)** lmmunofluorescence staining on paraffin embedded lung sections from Normal and CLAD lungs with α-sma antibody and **(C)** surfactant protein C antibody. Nuclei were stained with DAPI. N=3 Scale bar 100*µ*m **(D)** Quantification of fibrosis using Ashcroft score ***P<0.001 *t*-Test. **(E)** Quantification of number of SPC immunoreactive cells per high power field. Ten fields were used per group ****P<0.0001 *t*-Test. **(F-G)** Quantification of telomere length by calculating telomere fluorescence intensity over DAPI area in endobronchial and transbronchial biopsies diagnosed as CLAD and No CLAD **P<0.01 *t*-Test, *P<0.05 *t*-Test **(H-1)** Representative images of telomere quantitative fluorescence insitu hybridization (Q-FISH) assay conducted on endobronchial and transbronchial biopsies diagnosed with CLAD and No CLAD. **(J)** SA-β-gal staining (Blue colored cells) on OCT embedded cryosections. Counterstained with Eosin. N=6. **(K)** lmmunohistochemistry on paraffin embedded lung sections from normal and CLAD lungs with P16, P53 and P21 antibodies N=5.

### Telomere length and markers of senescence in CLAD lungs

It has been shown that telomeres are short in the proximal airway epithelium of patients at risk for developing CLAD (9). To determine whether telomeres are shortened in more than one epithelial cell subtype, endobronchial and transbronchial biopsies were used to measure telomere length by Telomere Q-FISH assay. Telomere lengths of patients with CLAD were significantly shorter in epithelial cells of endobronchial biopsies (Fig. 1F) identified by immunostaining with E-cadherin (Fig. 1H). In transbronchial biopsies, quantification of telomere length in SPC immunoreactive cells also revealed significantly shorter telomeres in CLAD lungs compared to lungs with no CLAD (Fig. 1 G and I).

Telomere dysfunction mediates cellular senescence reprogramming (21, 22), which is implicated as a driver of lung remodeling and fibrosis (15, 23). To examine whether senescence programming is activated in CLAD lungs, they were stained with X-gal chromogen to identify senescence-associated-Beta galactosidase (SA-β-gal) activity. Foci of SA-β-gal staining were found in CLAD lungs (Fig. 1J), indicating presence of senescent cells in epithelium lining the airway that appears to be obliterated. CLAD lungs were also immunostained for the senescence markers p16, p53, and p21 (Fig. 1J) revealing immunoreactive epithelial cells in regions of lung fibrosis.

### Telomere uncapping in club cells leads to short telomeres and increases mortality

Analysis of human samples demonstrated that telomere dysfunction and senescence programming are found in regions of lung remodeling and fibrosis observed in CLAD lungs. To test whether telomere dysfunction may mediate the remodeling, *TRF1^F/F^* mice were crossed with mice that inducibly express Cre recombinase in club cells upon treatment with tamoxifen (*SCGB1a1-Cre ERT2* mice) to generate *SCGB1a1-creTRF1^F/F^* mice. These mice were treated with tamoxifen weekly for up to 9-months (Fig. 2A). *SCGB1a1-creTRF1^F/F^* mice died between 7 to 9 months after beginning tamoxifen injections, in contrast to *TRF1^F/F^* mice, all of which survived (Fig. 2B). Telomere length was found to be approximately 4-fold shorter in club cells of *SCGB1a1-creTRF1^F/F^* mice compared to age-matched *TRF1^F/F^* controls (Fig. 2 C and D) when quantified by fluorescent *in situ* hybridization (Q-FISH).

**Fig. 2.**
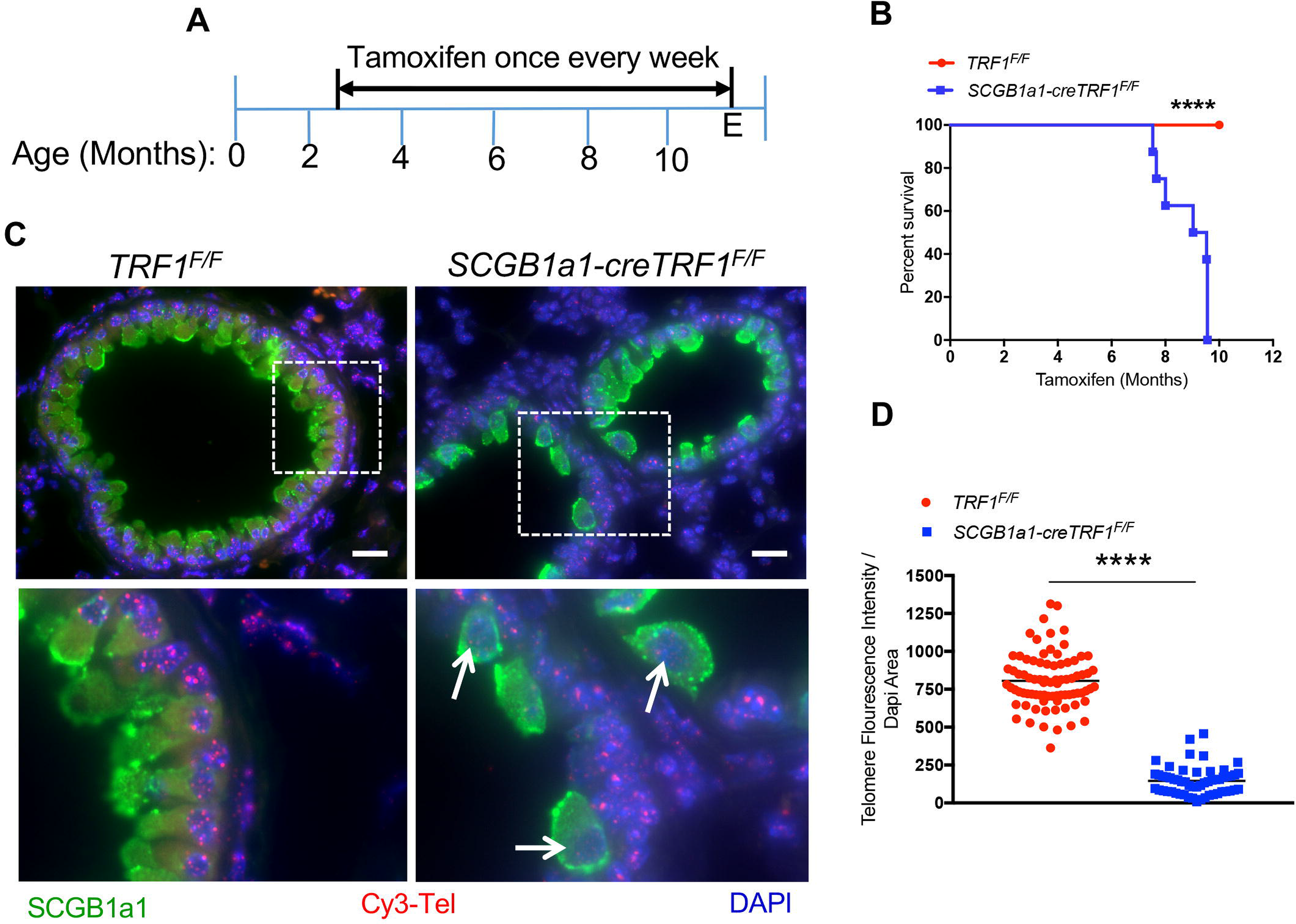
Deletion of TRF1 in club cells leads to short telomeres and increased mortality. **(A)** Tamoxifen-inducible ere-mediated recombination approach to delete TRF1 in ere expressed cells. Tamoxifen was administered to mice weekly starting from 10 weeks age intraperitoneally at 250 mg/kg body weight. E=End point **(B)** Kaplan-Meier Survival graph of *TRF1^F/F^* and *SCGB1a1-creTRF1^F/F^* mice treated with weekly injections of tamoxifen 250 mg/kg body weight. ****P<0.0001 *log-rank* test. **(C)** Telomere Quantitative fluorescence insitu hybridization (Q­ FISH) assay conducted on paraffin embedded lung sections collected after 8-9 months of tamoxifen administration. Cy3-Tel is PNA probe to detect TTAGGG repeats. SCGB1a1 antibody to detect club cells. DAPI to detect nuclei. Boxed area is shown in bottom panels. scale bar 20*µ*m. **(D)** Quantification of telomere length using telomere fluorescence intensity with reference to DAPI area N=3 mice/group. Each dot represents telomere fluorescence intensity in each club cell acquired from twenty different high power fields. Data is pooled from 3 mice. ****P<0.0001 *t*-Test.

### Telomere dysfunction in club cells leads to airway-centric lung remodeling and fibrosis

To evaluate the lung pathology of *SCGB1a1-creTRF1^F/F^*mice, lungs were stained with Masson’s trichrome stain. Increased collagen deposition was found predominantly around the small airways and extended into the adjacent lung parenchyma of *SCGB1a1-creTRF1^F/F^* lungs (Fig. 3A and Fig. S1A in supplemental data). Obliteration of small airways (Fig. S1B in supplemental data) and narrowing of the larger airway lumens was also observed (Fig. 3A and Fig. S1A in supplemental data). Ashcroft scores for fibrosis were consistently elevated in *SCGB1a1-creTRF1^F/F^*mice relative to controls (Fig. 3B). Quantification of collagen content in lungs by hydroxyproline assay revealed 3-fold more collagen in *SCGB1a1-creTRF1^F/F^*lungs (Fig. 3C) and qPCR of mouse lung mRNA demonstrated increased collagen 1 expression in *SCGB1a1-creTRF1^F/F^* lungs (Fig. 3D).

**Fig. 3.**
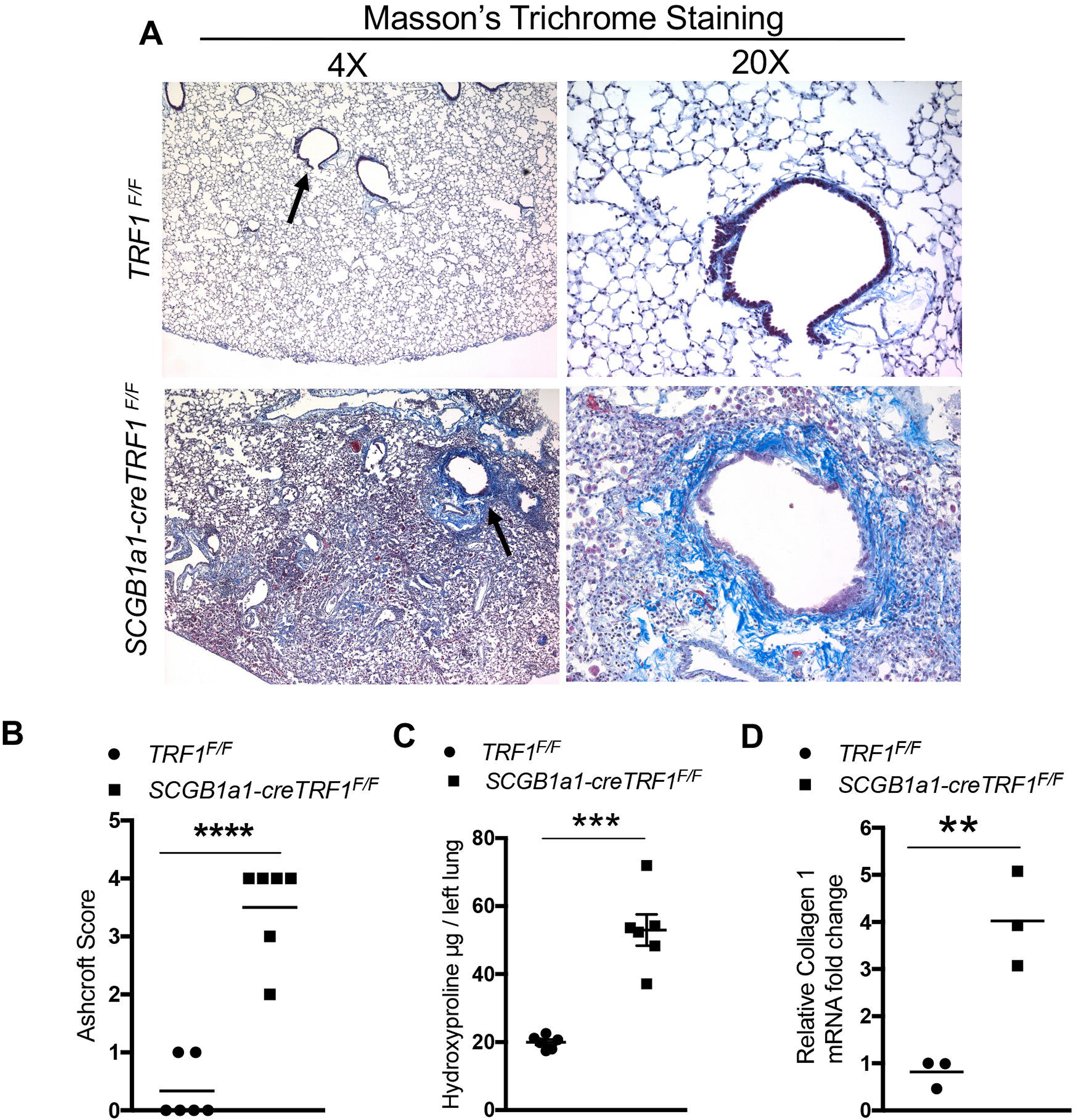
Airway-centric lung remodeling and fibrosis. **(A).** Masson’s Trichrome staining was performed on lung sections from *TRF1^F/F^* and *SCGB1a1-creTRF1^F/F^* mouse lungs treated with weekly doses of tamoxifen 250 mg/kg body weight. Deposition of collagen is indicated by blue stain. Magnification 4X and 20X. Arrows indicate areas of collagen deposition around bronchioles **(B)** Quantification of fibrosis using Ashcroft score ****P<0.0001 *t*-Test. **(C)** Quantification of collagen deposition by hydroxyproline assay on lung tissues from *TRF1^F/F^* and *SCGB1a1-creTRF1^F/F^* mice ***P<0.001 *t*-Test. **(D)** Quantitative PCR of collagen1 mRNA in lungs from *TRF1^F/F1^* and *SCGB1a1-creTRF1^F/F^* mice 9 months after tamoxifen administration **P<0.001 *t*-Test.

To assess overall lung morphometry, micro-computed tomography was performed on lungs extracted from *SCGB1a1-creTRF1^F/F^* and *TRF1^F/F^*control mice 9 months after tamoxifen administration. Coronal sections demonstrated increased density of tissue in central regions of *SCGB1a1-creTRF1^F/F^* lung consistent with bronchiolar fibrosis (Fig. 4A). Cross-sectional images show decreased airway frequency and diameter in *SCGB1a1-creTRF1^F/F^*lungs compared to control lungs (Fig. 4A).

**Fig. 4.**
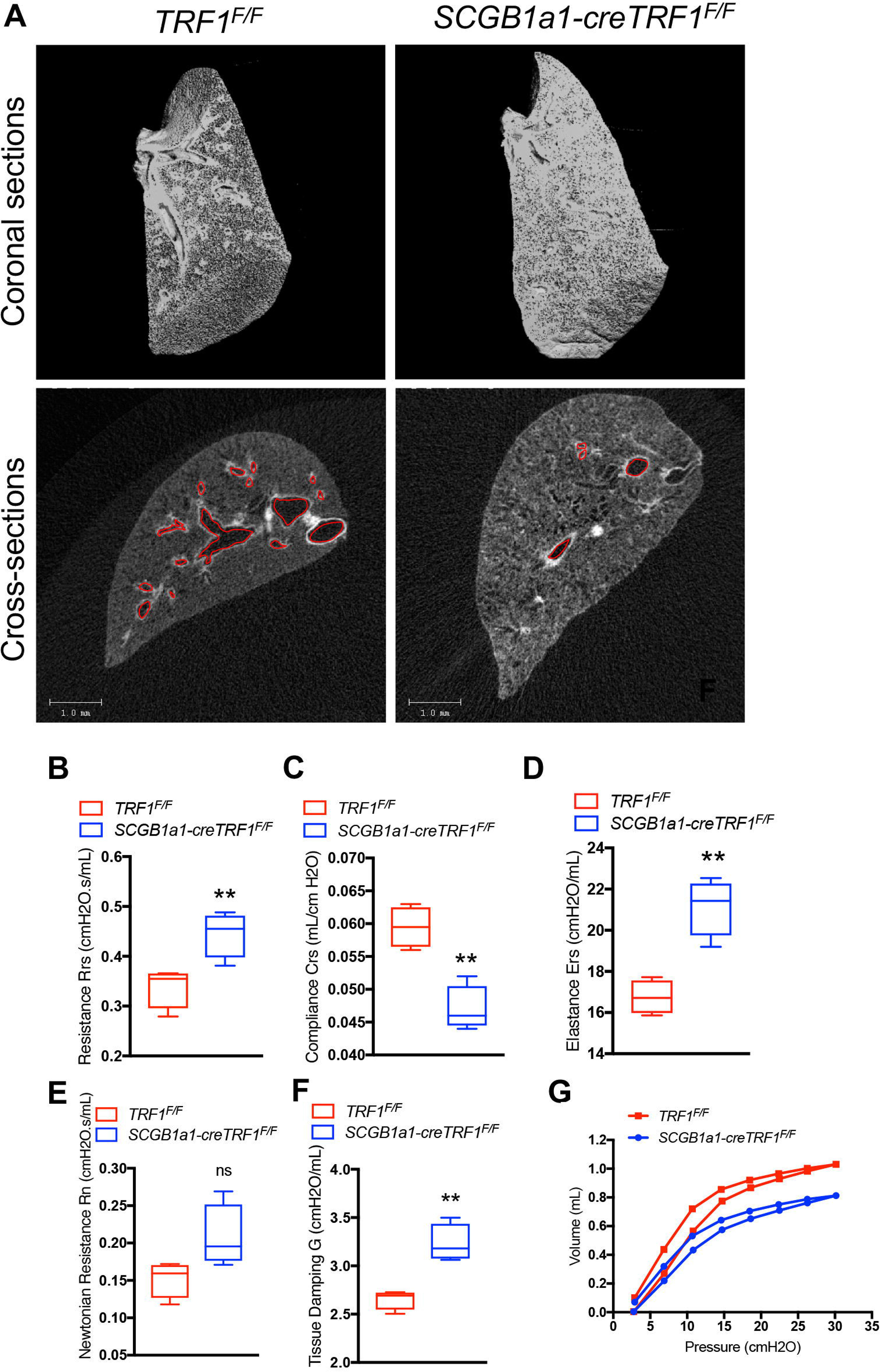
microCT imaging and lung mechanics. **(A)** Lungs harvested from *TRF1^F/F^* and *SCGB1a1-creTRF1^F/F^* mice following treatment for 9 months with weekly injections of tamoxifen (250 mg/kg body weight) were subjected to microCT imaging. Left lungs were used for imaging. Top panel: Coronal section passing through the middle of the lung visualized in 3D. Air spaces are represented in black. Grey represents tissue density. Bottom panel: Cross-sectional images showing air-spaces, airways and tissue density. Airways are marked with red border. Scale bar 1mm. **(B-G)** Lung mechanics measured by flexivent. *TRF1^F/F^* and *SCGB1a1-creTRF1^F/F^* mice after treatment for 9 months with weekly injections of tamoxifen were subjected to measurement of total resistance (Rrs), Compliance (Crs), Elastance (Ers) Airway resistance (Rn), tissue damping (G) and Pressure-Volume (PV) loops. N = 4 mice/group **P < 0.01. *t*-test ns=not significant.

### Lung function of *SCGB1a1-creTRF1^F/F^* mice

As CLAD is defined in part by obstructive airway physiology, we evaluated the impact of club cell telomere dysfunction on lung function of *SCGB1a1-creTRF1^F/F^*compared to *TRF1^F/F^* control mice using the flexivent instrument. Resistance of the overall respiratory system (Rrs) (Fig. 4B) was higher in *SCGB1a1-creTRF1^F/F^* lungs compared to control lungs. Lung compliance (Crs) (Fig. 4C) was significantly lower with an analogous increase in elastance (Fig. 4D) of the respiratory system (Ers) in *SCGB1a1-creTRF1^F/F^* lungs. To differentiate lung mechanics of central and peripheral lung compartments, broadband forced oscillations (constant phase model) were used. In this model, newtonian resistance (Rn) and tissue damping (G) were increased in *SCGB1a1-creTRF1^F/F^* lungs relative to controls (Fig 4, E and F). We did not observe a statistically significant difference in newtonian resistance (Rn), a parameter indicative of large conducting airways resistance (Fig. 4E). Overall, this pattern of abnormality suggests increased resistance of small airways in the peripheral compartment and increased alveolar tissue stiffness. Quasi-static lung mechanics measured using Pressure-volume (P-V) curves (Fig. 4G) showed that the static compliance (Cst), a parameter which assesses lung distensibility, was significantly reduced in *SCGB1a1-creTRF1^F/F^* lungs compared to *TRF1^F/F^* control lungs. In comparison, lung mechanics of *SPC-cre TRF1^F/F^*mice show similar trend as *SCGB1a1-creTRF1^F/F^* lungs except that tissue damping (*G*) is not significantly different compared to *TRF1^F/F^* control lungs, confirming that distal airways are not remodeled and the location of fibrosis is spatially different in *SPC-cre TRF1^F/F^* mice (Fig. S2 in supplemental data).

### Immune response profile in *SCGB1a1-cre TRF1^F/F^* mice

Bronchoalveolar lavage (BAL) fluid collected from *SCGB1a1-cre TRF1^F/F^* mice showed elevated levels of the pro-fibrotic marker TGF-β1 (Fig. 5A), senescence-associated cytokine IL-6 (Fig. 5B) and chemokine CCL2 (Fig. 5C), compared to *TRF1^F/F^* mice. Analysis of BAL fluid revealed greater than 6-fold increase in total cell count in *SCGB1a1-cre TRF1^F/F^* mice compared to *TRF1^F/F^* mice (Fig. 5D). Differential cell count of BAL fluid indicated a significant increase in polymorphonuclear (PMN) granulocytes (Fig. 5E), macrophages (Fig. 5F), and lymphocytes (Fig. 5G). Immunophenotyping of whole lung homogenates revealed increased CD45+ leukocytes, including both T and NK cells (Fig. 5H and Table S1 in supplemental data) in *SCGB1a1-cre TRF1^F/F^* mice compared to *TRF1^F/F^* mice.

**Fig. 5.**
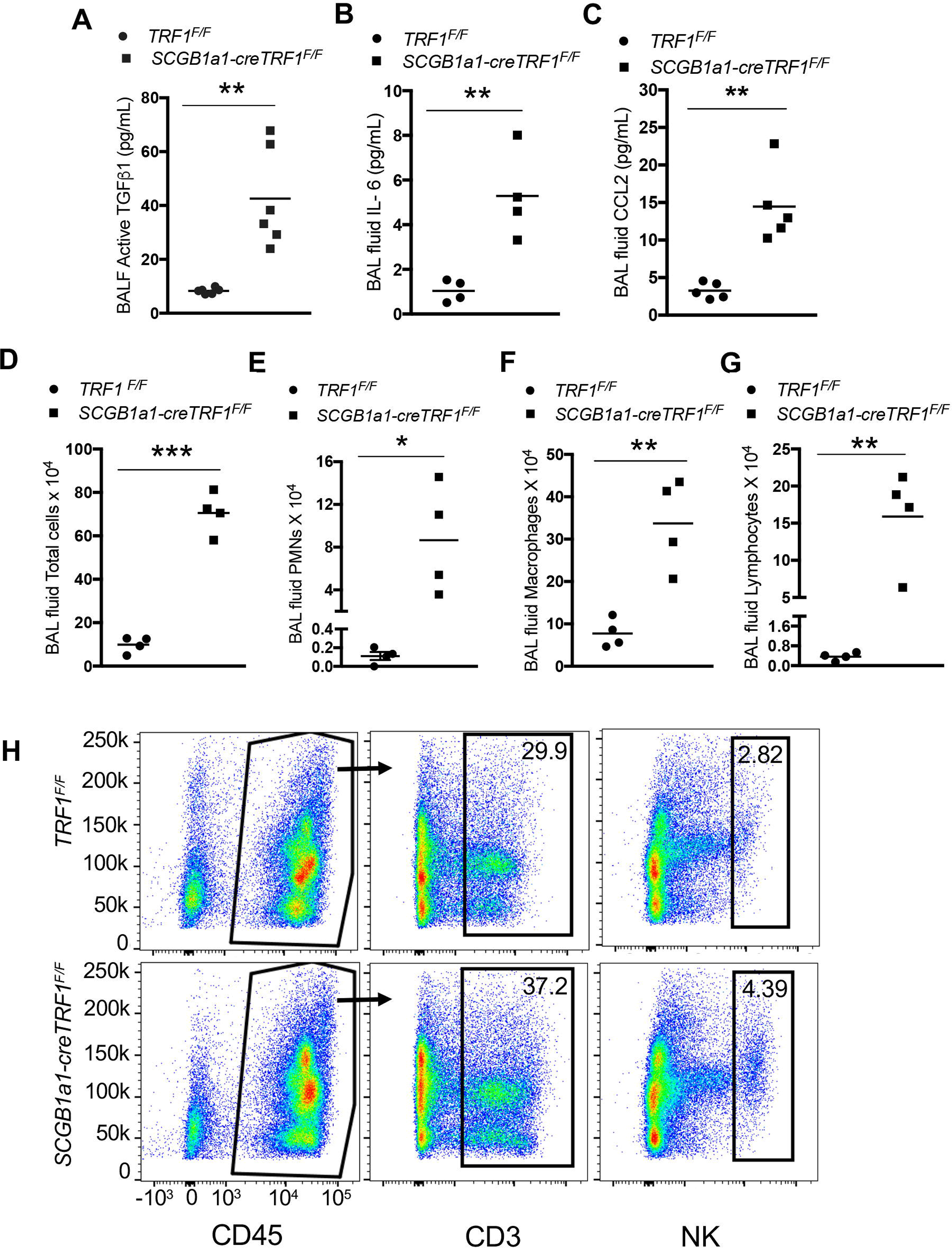
Mouse BAL fluid ELISA, cell count differential and lung immune cell analysis. **(A-C)** ELISA on BAL samples from *TRF1^F/F^* and *SCGB1a1-creTRF1^F/F^* mice to test for Active TGF-B1, IL-6 and CCL2. **(D-G)** Cell count and differentials of BALF samples from lungs of *TRF1^F/F^* and *SCGB1a1-creTRF1^F/F^* mouse lungs treated with weekly doses of tamoxifen for 9 months. N=4 mice/group. *P<0.05, **P<0.01, ***P<0.001 *t*-test. **(D)** Total cell count. **(E)** PMNS **(F)** Macrophages and **(G)** Lymphocytes **(H)** lmmunophenotyping of lungs from *TRF1^F/F^* and *SCGB1a1-creTRF1^F/F^* at 9 months by flow cytometry. Percentage of CD3 and NK cell populations were derived from CD45 positive gate. N=3. Individual population numbers are mentioned in Supplementary Table 1.

### TRF1 deletion in club cells leads to DNA damage and activation of cellular senescence programming

To determine the consequence of telomere dysfunction within club cells, immunostaining was carried out for the DNA damage marker gamma-H2AX. DNA damage foci localized to club cells (Fig. 6A) of *SCGB1a1-cre TRF1^F/F^* lungs. Moreover, SA-βgal staining to identify senescent cells showed blue stained cells in the airway epithelium in *SCGB1a1-cre TRF1^F/F^* lungs compared to *TRF1^F/F^* controls, consistent with activation of senescent programming (Fig. 6B).

**Fig. 6.**
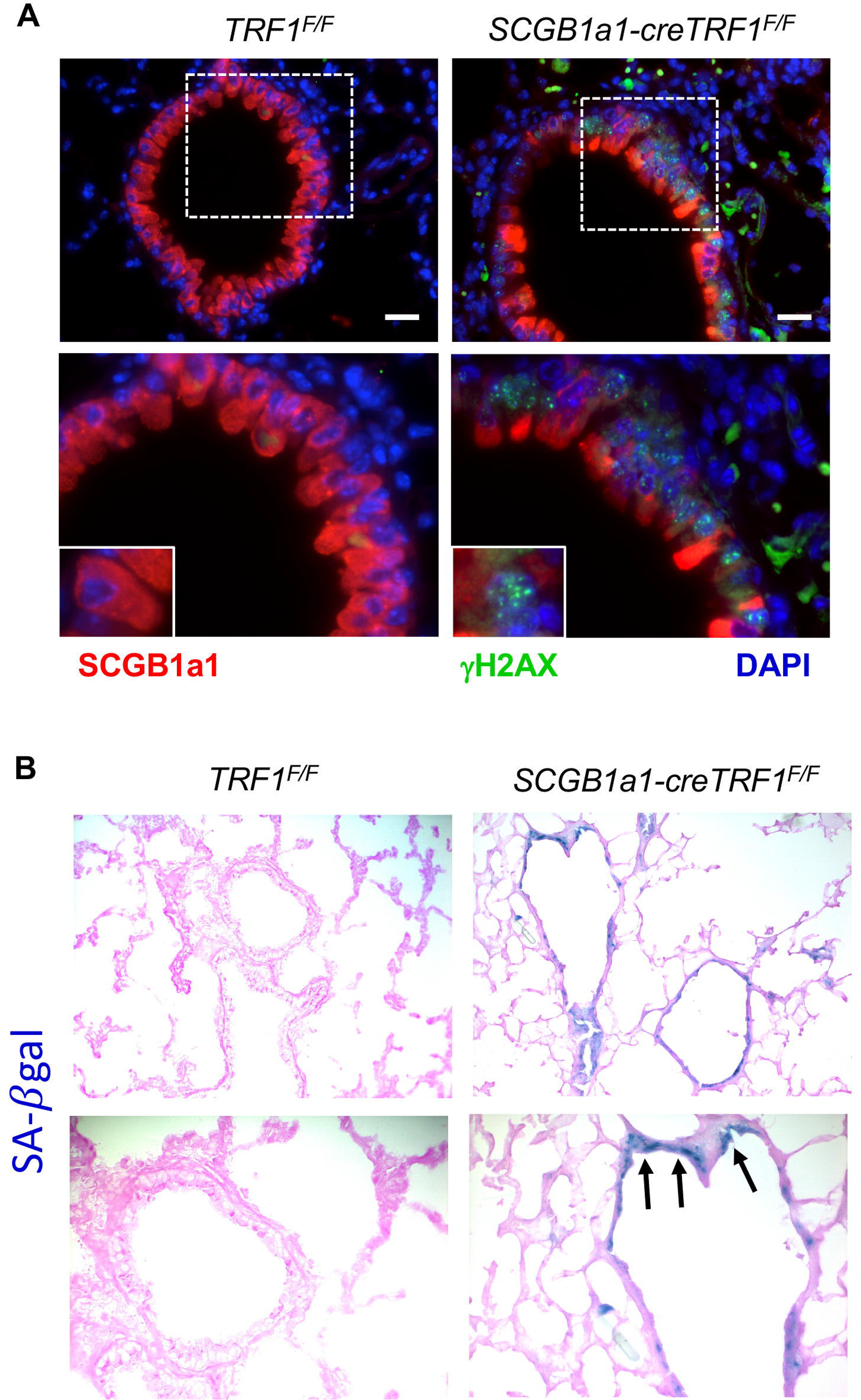
Immunostaining of lung sections and detection of cellular senescence by SA-*β* galactosidase activity. **(A)** Immunostaining on sections of lungs harvested from *TRF1^F/F^* and *SCGB1a1-creTRF1^F/F^* mice treated with weekly tamoxifen injections for 9 months. **(A)** SCGB1a1 immunostain for club cells (red) and *γ*H2AX (green). *γ*H2AX foci (green) in the nuclei of club cells. Nuclei were stained with DAPI (blue). Scale bar 20*µ*m Boxed area is enlarged in bottom panel along with inset showing individual club cell **(B)** SA-β-gal staining of sections of lungs harvested from *TRF1^F/F^* and *SCGB1a1-creTRF1^F/F^* mice following treatment for 9 months with weekly injections of tamoxifen (250 mg/kg body weight). Note the blue SA-*β*gal+ cells (arrows) in lungs of *SCGB1a1-creTRF1^F/F^* mice.

### Club cell telomere dysfunction leads to loss of club cells and proliferation of type II AECs

Immunofluorescent co-staining for club cells (*SCGB1a1*), type II AECs (SPC), and the cell proliferation marker Ki-67 showed fewer *SCGB1a1* immunoreactive cells and increased expression of SPC and Ki67 in *SCGB1a1-cre TRF1^F/F^*lungs compared to *TRF1^F/F^* controls (Fig. 7A). Quantification of cell numbers revealed a 5-fold increase in total Ki67+ cells (Fig. 7B), a 2-fold increase in SPC+ cells (Fig. 7C) and 8-fold increase in dual SPC+Ki67+ cells (Fig. 7D). in *SCGB1a1-cre TRF1^F/F^* lungs compared to *TRF1^F/F^* controls. Within SPC cells, there was a 4-fold increase in Ki67 positivity in *SCGB1a1-cre TRF1^F/F^* lungs over controls (Fig. 7E). These results demonstrate a loss of *SCGB1a1* immunoreactivity, type II cell hyperplasia, and increased proliferation of type II AECs. Moreover, quantitative PCR of lung lysates from *SCGB1a1-cre TRF1^F/F^*mice showed more than 5-fold increase in SMA mRNA expression levels, ∼1.5-fold increase in SPC and nearly a 3-fold decrease in *SCGB1a1*expression compared to *TRF1^F/F^* mice (Fig. 7F). Immunofluorescence staining demonstrated increased SMA expression around airways (Fig. 7G).

**Fig. 7.**
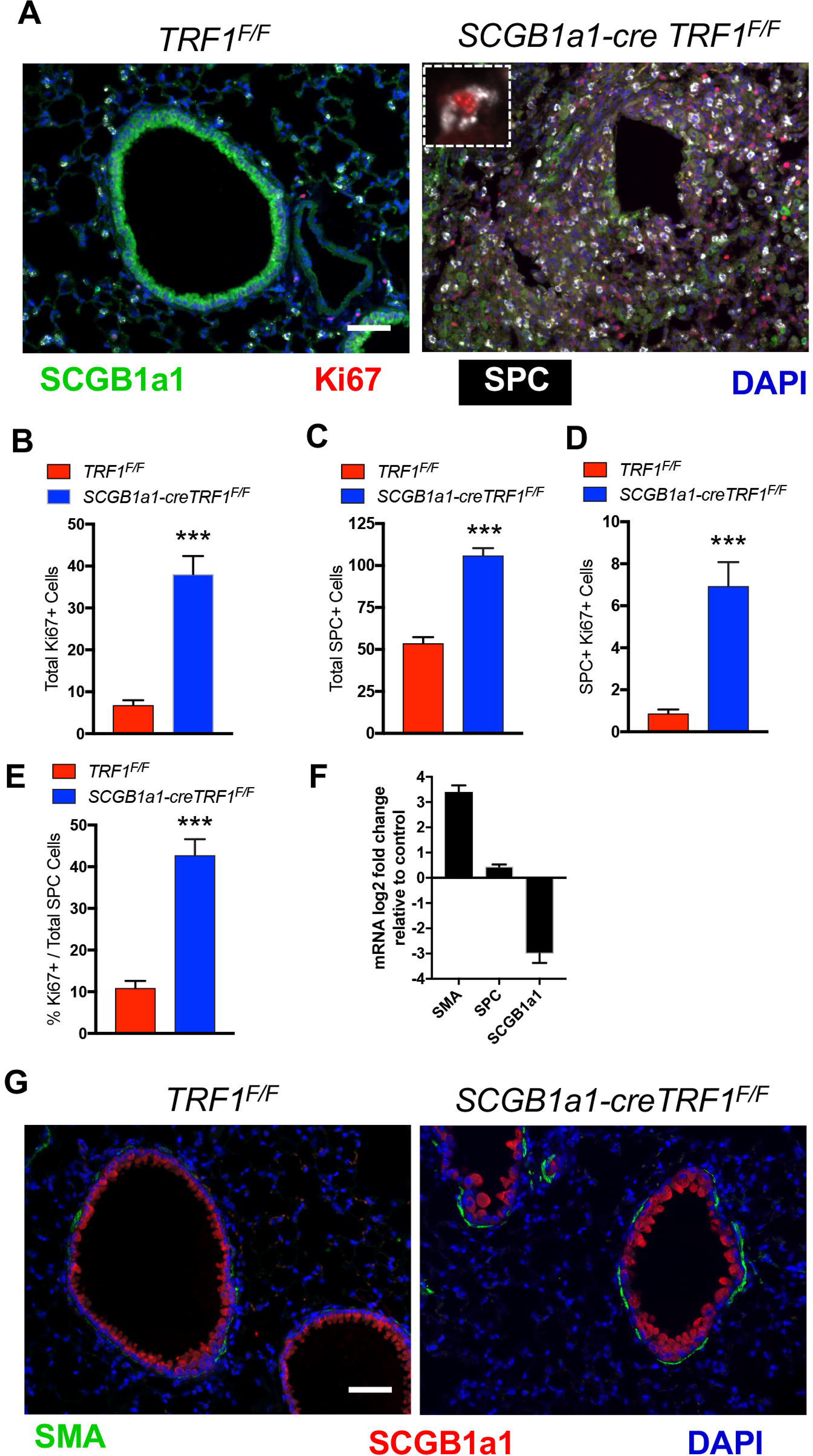
Loss of club cells and type II AEC hyperplasia around bronchioles. **(A)** Immunostaining on lung sections from *TRF1^F/F^* and *SCGB1a1-creTRF1^F/F^* mice treated with weekly doses of tamoxifen. Cell proliferation marker Ki67 (red), SCGB1a1 (green) for club cells and SPC (Far red changed to white for visibility). *Inset:* Ki67 marker in an SPC+ cell. Nuclei were stained with DAPI (blue) Scale bar 50*µ*m. **(B)** Quantification of total Ki67 positive cells, **(C)** total SPC+ cells **(D)** SPC and Ki67 double positive cells **(E)** Percentage ki67+ cells over total SPC positive cells **(F)** mRNA expression levels in *SCGB1a1-creTRF1^F/F^* lungs represented as log2 fold change over *TRF1^F/F^* controls **(G)** lmmunofluorescence staining on paraffin embedded lung sections co-stained with alpha-SMA and SCGB1a1. Scale bar 50*µ*m.

## DISCUSSION

This manuscript reports that CLAD lungs have relatively short telomeres and express senescence markers in airway epithelial cells. Modeling this telomere dysfunction in club cells resulted in physiologic and pathologic changes that are similar to those found in patients with CLAD (Table S2 in supplemental data). These changes included airway obstruction, with airway-centric fibrosis and obliteration of bronchioles; parenchymal fibrosis; expansion of type II AEC cell numbers adjacent to the obliterated bronchioles; and regional lung fibrosis (24). Consistent with obliterated airways, the lung physiology of *SCGB1a1-cre TRF1^F/F^*lungs demonstrated increased tissue damping (*G*) reflecting increased peripheral airways resistance (25). Elements of CLAD pathology were recapitulated by modeling a single molecular defect in the absence of alloimmunity. Collectively, these data suggest that the airway epithelial cell telomere dysfunction is a molecular driver of CLAD pathology.

Short telomeres are evident in diseases of tissue remodeling and fibrosis including skin dyskeratosis (26), hypertrophic cardiomyopathy (27–29), liver cirrhosis (30), Duchenne muscular dystrophy (31), idiopathic pulmonary fibrosis (32), and CLAD (9). Telomere dysfunction secondary to deletion of shelterin protein TRF1 in specific cell types has modeled organ-specific pathologic changes attributed to human syndromes of telomere dysfunction. For example, deletion of TRF1 in keratinocytes or hematopoietic cells caused hyperpigmentation and bone marrow failure, respectively (12, 13). In the lung, deletion of TRF1 in type II AECs leads to lung fibrosis in a subpleural distribution (15). In distinction with the mouse model described here, lungs of animals with type II AEC telomere dysfunction showed no difference in peripheral airway resistance (*G)* values when compared to controls, suggesting that physiological and pathological outcomes depend on the cell type where telomeres are dysfunctional. Overall, it appears that telomere dysfunction may be a molecular driver of many forms of tissue remodeling, but the specific pathologic appearance of the remodeling depends on the tissue and cellular subtype within the tissue where the telomere dysfunction is found.

Telomere shortening and uncapping activates DNA damage machinery, cell cycle checkpoints, and senescence reprogramming (21, 22). Consistent with these reports, we found γH2AX-positive DNA damage foci and positive staining for cellular senescence markers in *SCGB1a1-cre TRF1^F/F^* transgenic mice, which could amplify local inflammatory responses (33). Accumulation of senescent cells correlated temporally to the development of fibrosis suggesting their contribution to fibrosis. Critically short telomeres in club cells causing cellular senescence may lead to release of senescence-associated secretory proteins (SASP) such as IL-6 or CCL2 into the local lung mileu leading to fibrotic remodeling around the airways.

Club cells are primarily located in the trachea and terminal bronchioles where they function as stem cells by differentiating into other epithelial cell subtypes (34). In response to lung injury, club cells are shown to give rise to type I and type II alveolar epithelial cells (35). Telomere shortening was associated with the loss of cells expressing *SCGB1a1* and increased Ki67 staining in SPC immunoreactive cells adjacent to the airways. This raises the possibility that type II AECs expand to compensate for the loss of functional club cells. An alternative explanation is that short telomeres and senescence reprogramming in club cells leads to their de-differentiation, diminishing their ability to express *SCGB1a1* and hence they could go undetected when probed. Overall, abnormal regeneration of lung epithelium secondary to club cell dysfunction may represent a key pathway for the propagation of CLAD that would be relatively resistant to immunosuppression. Although, it is not known to what extent the observed pathology is attributable to club cell telomere dysfunction, as opposed to club cell injury in general, the observations of senescence, type II AEC proliferation, and subsequent leukocyte recruitment all suggest a specific role for telomere dysfunction.

While multiple immune mechanisms likely contribute to CLAD, this syndrome is generally linked to alloimmunity (36). Interestingly, the increased numbers of parenchymal and BAL immune cells observed in *SCGB1a1-cre TRF1^F/F^* transgenic mice revealed an immune response in the lung consistent with those found in CLAD lungs, however in the absence of alloimmunity. We speculate that senescence reprogramming due to telomere dysfunction in club cells leads to release of an array of cytokines and chemokines that attract immune cells to the lung. Elevated numbers of T cells and NK cells could reflect a direct response to cell stress (37) or be due to T-cell attracting chemokines released by senescent club cells, such as CXCL-9, CXCL-10 and CXCL-11, that are commonly elevated in airway inflammation (38) and CLAD (39, 40). Elevated levels of lymphocytes in lungs of *SCGB1a1-cre TRF1^F/F^* transgenic mice suggests that telomere dysfunction in club cells could even contribute to the inflammatory changes seen in CLAD. These findings have important implications for immunosuppression strategies in lung transplantation, as targeting alloreactive T cells might not alter the trajectory of a disease driven by club cell telomere dysfunction.

Currently available mouse models of lung transplantation (41) have certain limitations. Animal models of tracheal transplantation (42–45) reproduce airway luminal narrowing, but may not reflect the small airway disease, parenchymal changes, and vascularity of human allografts. MHC mismatched orthotopic transplantation models reproduce features of acute rejection, but not necessarily CLAD. (46–49). Transplantation of B6D2F1/J lungs to DBA recipients has shown promise in development of CLAD-like pathology, but the DBA background may be a limitation because of the paucity of available genetic knockout models. Telomere dysfunction is a molecular defect identified in CLAD. Remarkably, the mouse model presented in this manuscript recapitulates changes found in CLAD by genetic modification of telomere protein. Modeling the elements of CLAD pathology mediated by telomere dysfunction in airway epithelial cells has the advantage that surgery is not required. At the same time, since *SCGB1a1-cre TRF1^F/F^* transgenic mice model a single molecular pathway resulting in CLAD pathophysiology, this model may be less informative regarding other important mechanisms relevant to lung transplantation, including ischemia reperfusion injury, revascularization, alloimmunity, and immunosuppression.

In summary, telomere dysfunction in club cells of *SCGB1a1-cre TRF1^F/F^* transgenic mice causes progressive lung remodeling and airway-centric fibrosis, similar to the lung remodeling and fibrosis characteristic of CLAD. The findings reported in this manuscript support the hypothesis that telomere dysfunction is an alloimmune independent driver of the lung remodeling and fibrosis found in patients with CLAD. In the context of lung transplantation, the lung epithelium undergoes high rates of epithelial cell turnover, this may lead to accelerated telomere attrition, senescence reprogramming of airway progenitor cells and ultimately airway remodeling and fibrosis. The findings also suggest that CLAD and other forms of age-related lung remodeling such as IPF may share common molecular drivers. The *SCGB1a1-cre TRF1^F/F^* spontaneous mouse model of lung remodeling may be useful for studying the contribution of molecular mechanisms of lung remodeling, driven by short telomeres in airway epithelial cells, to the airway-centric fibrosis observed in CLAD.

## Acknowledgments

We are grateful to all the patients and coordinators who participated in this study. We thank Alfred Li at the Bone Imaging Research Core, UCSF for assistance with microCT imaging.

## Data and materials availability

All data related to this study are present in the paper or the supplementary materials

## Author contributions

R.P.N. and P.J.W. designed the study and drafted the manuscript. R.P.N. performed experiments, analyzed and interpreted data and wrote the manuscript. G.G., N.A.Z., and J.E.M. performed experiments and analyzed data. K.D.J., J.K., and J.R.G. contributed to study design, data interpretation, and manuscript revision. All authors reviewed and approved of the final manuscript.

