## Supplemental materials for "Telomere Dysfunction Drives Chronic Lung Allograft Dysfunction Pathology"

Supplemental Material

Fig. S1

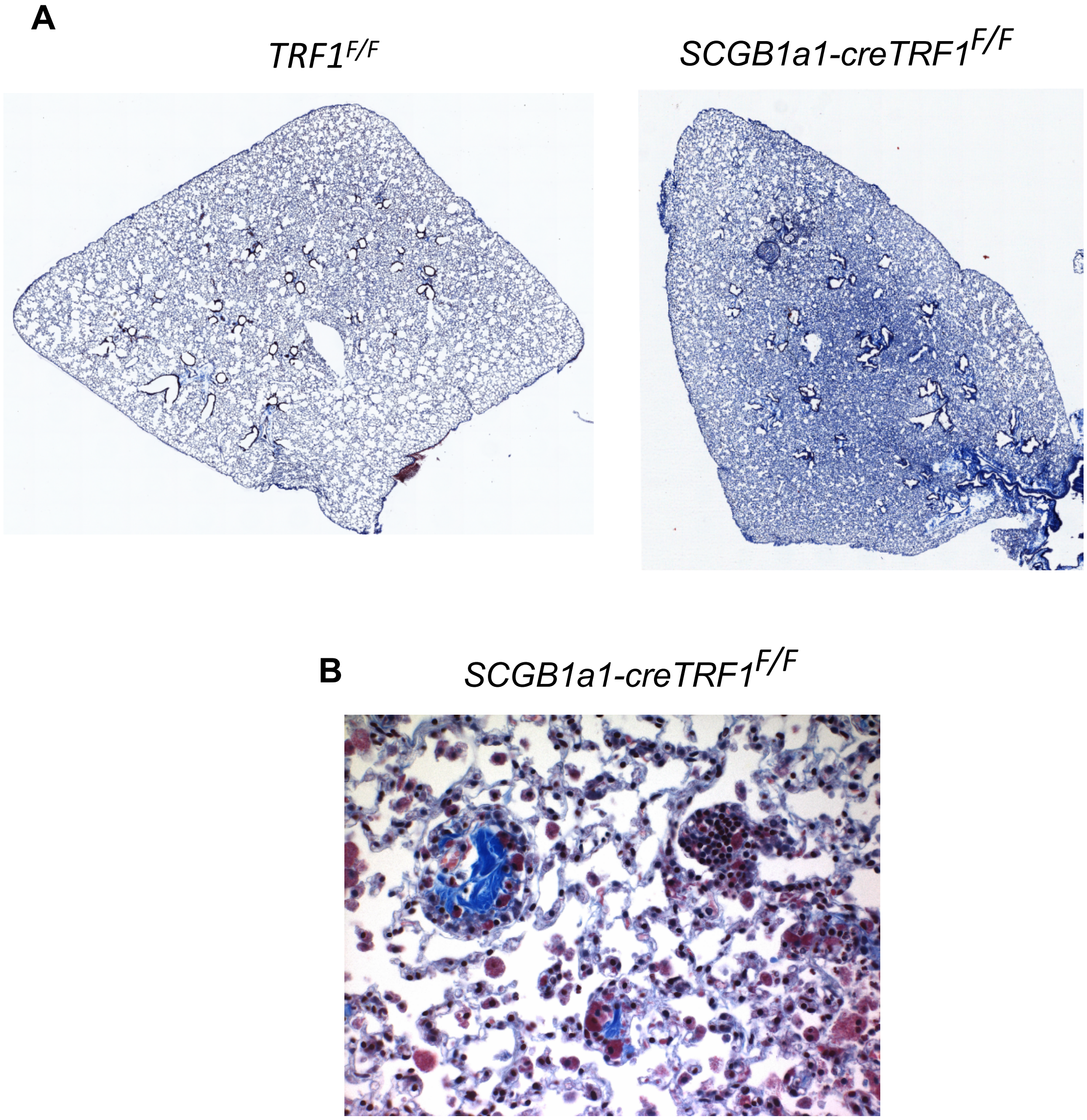


**Fig. S1. Airway-centric lung remodeling in *SCGB1a1-creTRF1$^{F/F}$* mice (A)** Mosaic images of Masson's trichrome stained mouse lungs from *TRF1$^{F/F}$* and *SCGB1a1-creTRF1$^{F/F}$* mice treated with weekly doses of tamoxifen for 8 months. (B) Lungs from *SCGB1a1-creTRF1$^{F/F}$* showing obliteration of small airways with collagen (blue).

Fig. S2

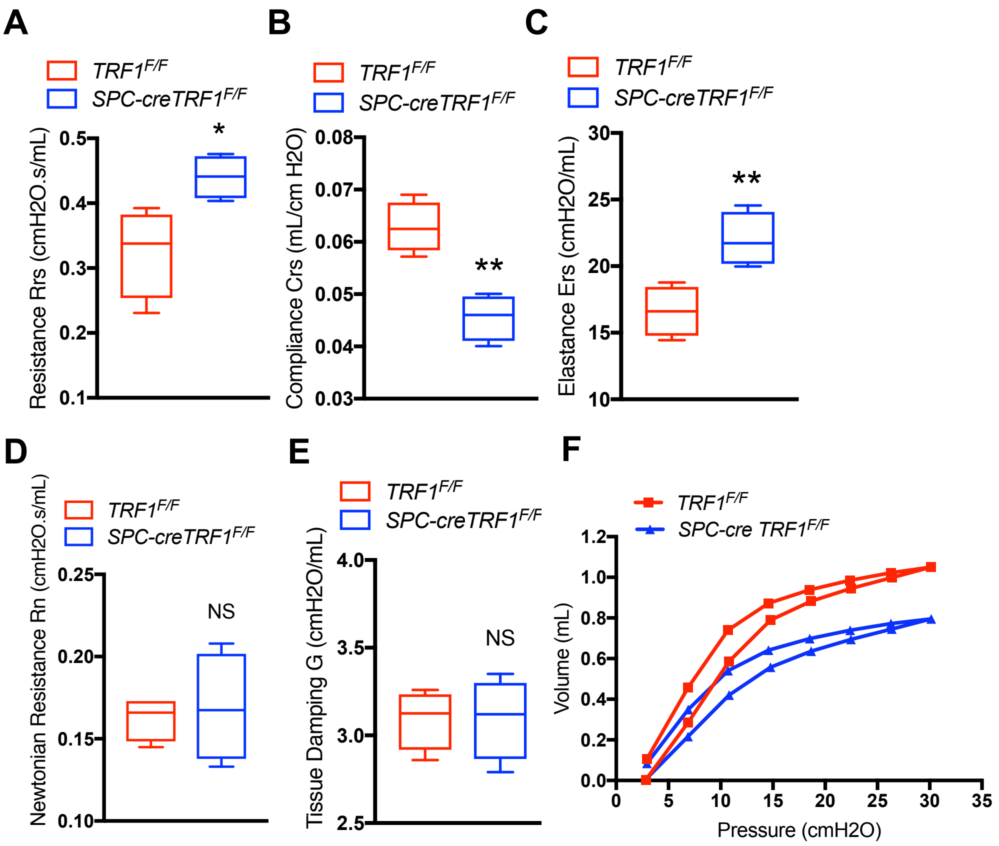


**Fig. S2. Lung mechanics measured by flexivent in SPC-cre transgenic mice** Lung mechanics were measured by flexivent in *TRF1*$^{F/F}$ and *SPC-creTRF1*$^{F/F}$ mice following treatment for 9 months with weekly injections of tamoxifen (250 mg/kg body weight). Graphs **(A-C)** represent measurement of total resistance (Rrs), Compliance (Crs), Elastance (Ers) respectively. **(D-E)** Graphs represent Newtonian resistance (Rn) and tissue damping (G) respectively. **(F)** Graph represents pressure-volume loops. N = 4 mice/group *P < 0.05; **P < 0.01. *t*-test ns=not significant.

### Table S1

| Cell type | *TRF1 F/F* | *SCGB1a1-cre TRF1 F/F* |
|---|---|---|
| | Cell number (x10$^4$) Mean ± SEM | |
| CD45 positive (Leukocytes) | 514.9 ± 75.66 | 848.6 ± 55.49 |
| CD3 positive (T-cells) | 176.5 ± 45.65 | 333.8 ± 29.37 |
| NK cells | 31.25 ± 18.89 | 116.6 ± 50.17 |

**Table S1. Quantification of cell numbers from immunophenotyping mouse lungs by flow cytometry:** *TRF1$^{F/F}$* and *SCGB1a1-creTRF1$^{F/F}$* mouse lung obtained after 9 months of tamoxifen administration were analyzed by flow cytometry. Table shows individual cell populations obtained after gating whole population on Live/Dead, CD45, CD3 and NK. Individual populations were derived by multiplying percentages obtained with total cell populations in lung homogenate from respective groups. N=3

| Feature | CLAD | Mouse model |
|---|---|---|
| Airway-centric lung fibrosis | Yes | Yes |
| Type II cell hyperplasia | Yes | Yes |
| Short telomeres | Yes | Yes |
| SMA deposition | Yes | Yes |
| Cellular Senescence | Yes | Yes |
| Loss of Club cells | Yes | Yes |
| Loss of airways | Yes | Yes |
| Bronchiolitis obliterans | Yes | Yes |
| Compromised lung function | Yes | Yes |
| Pleuroparenchymal Fibroelastosis (PPFE) | Yes | No |

**Table S2. Comparison of features in CLAD vs mouse model:** Table shows similarities and differences between CLAD vs mouse model of telomere dysfunction in *SCGB1a1-creTRF1*$^{F/F}$ mice. Features listed signify the type of fibrosis, cell type, location, cellular and molecular features.

#### MATERIALS AND METHODS

**Study design:** The purpose of this study was to test the hypothesis whether short telomeres are cause or consequence of chronic lung allograft dysfunction. To test this hypothesis both human samples and C57BL/6 mice were used. Sample size for each experiment is indicated in the figure legend. Animal sample sizes were based on pilot experiments. End points of animals was based on pilot studies. Animals were randomly assigned to groups. Data collection and analysis were performed in a blinded manner. No outliers were excluded from the study.

**Human specimens:** Fibrotic lung tissue was obtained at the time of lung transplantation from patients with a pathological diagnosis of chronic lung allograft dysfunction. Control human lungs consisted of lungs not used by the Northern California Transplant Donor Network; our studies indicate that these lungs are physiologically and pathologically normal (1). Endobronchial and transbronchial biopsies were performed within 90 days post-transplant and sections were recut from FFPE tissue blocks, as previously described (2). Written informed consent was obtained from all subjects and was approved by the University of California, San Francisco Institutional Review Board protocol number 13-10738.

**Mice:** *TRF1*$^{F/F}$ mice were purchased from Jackson lab, *SCGB1a1-cre/ERT* mice were provided by Jason Rock, at UCSF. Mice were bred and housed in pathogen-free conditions in accordance with the guidelines of Laboratory Animal Resource Center (LARC). All animal procedures were carried out using protocols approved by the Institutional Animal Care and Use Committee (IACUC) at the University of California, San Francisco.

**Tamoxifen administration:** Tamoxifen (Toronto Research Chemicals T0600), suspended in peanut oil was injected (250 mg/kg body weight once per week) intraperitoneally to *SCGB-creTRF1*$^{F/F}$ mice and *TRF1*$^{F/F}$ littermate controls beginning at 10 weeks age.

**Hydroxyproline assay:** Hydroxyproline levels were quantified as described previously (3). Murine lungs were homogenized in 1 ml water. One hundred and twenty-five microliters of 50 % Trichloroacetic acid was added and the suspension incubated on ice for 20 min. Samples were centrifuged at 800 *g* for 5 min at 4ºC. The supernatant was discarded and 1 ml of 12 N HCl added to the pellet, which was then baked in an oven at 110ºC for 24 h. Dried pellets were reconstituted in 2 ml distilled water. Sample or serially diluted 6-Hydroxyproline standard (200 µl) was added to Chloramine T solution (1.4% chloramine T in 0.5 M Na Acetate and 10% isopropanol) and incubated at room temperature for 20 min. Five hundred microliters of Ehrlich's solution (1 M p-dimethylaminobenzaldehyde in 70% isopropanol and 30% perchloric acid) was added and the mixture incubated at 65ºC for 15 min and absorbance measured on a spectrophotometer at 550 nm.

**Histopathology and Immunofluorescence Staining:** Lungs were perfused with 10 % formalin at 20 cm $H_2O$ and fixed overnight before embedding in paraffin and sectioning at 4 µm thickness. For histopathology, tissues were subjected to hematoxylin & eosin staining or Masson's trichrome staining. For immunohistochemistry, tissues were deparaffinized in Xylene twice for 5 min each, subjected to ethanol gradient (100%, 95%, 75% and 50%) 3 min each followed by hydration for 2 min. Antigen retrieval was performed by heating sections to 95ºC for 20 min in citrate buffer (pH 6.0) and allowed to cool for 30 min. After PBS wash for 3 times, tissue sections were permeabilized with 0.1 % triton X-100 and Sections were blocked with 3 % BSA in donkey serum or goat serum and PBS) or MOM blocking kit (Vector labs BMK 2202) where applicable and incubated with primary antibodies SPC (sc-7706), SMA (Dako Cytomation M0851), phospho-γH2AX (ab26350), SCGB1a1 (sc-390313; sc-25555), p16 (sc-65224), p21 (sc-6246), p53 (sc-263), Ki67 (ab15580), E-cadherin (Cell signaling 3195S), pro-SPC (ab3786) overnight at 4ºC. Tissues were washed thrice in PBS tween and incubated with unconjugated secondary antibodies goat anti-mouse IgG1 (sc-2060), goat anti-mouse IgG2B(ab98703), goat anti-mouse IgG2A (ab97245) for DAB staining and fluorescent conjugated secondary antibodies donkey anti-mouse AlexaFluor 488, 546, 594 and 647 for fluorescent immunostaining at room temperature for 1 h followed by 3 washes. For DAB stained sections, slides were mounted using cytoseal whereas prolong gold anti-fade mounting medium with DAPI (Life Technologies) was used for fluorescent labeling. Images were acquired on Zeiss Axio Imager 2 microscope (Zeiss, Oberkochen, Germany).

**Bronchoalveolar lavage fluid (BALF) collection:** Bronchoalveolar fluid was collected from euthanized mice by lavaging lungs with 1 ml PBS. Recovered BAL fluid was centrifuged at 800 *g* for 5 min at 4ºC and the supernatant recovered for analysis. Red blood cells were lysed in the cell pellet with ammonium-chloride-potassium (ACK) buffer, the cells re-centrifuged and then suspended in 1 ml of PBS. The total numbers of cells were counted using a hemocytometer. Cells were cytospun and cell differentials determined by modified Giemsa stain (Electron Microscopy Sciences, Hatfield, PA).

**Telomere Q-FISH assay:** Telomere lengths were measured on paraffin-embedded tissue sections by Quantitative Fluorescence in Situ Hybridization (Q-FISH). Briefly, after deparaffinization, tissues underwent antigen retrieval by heating in microwave in 10mM sodium citrate buffer, pH

6.0, then incubating for 15 min in 0.01 M HCL containing 1% pepsin (Thermofisher Scientific, South San Francisco, CA). Tissues were washed and air-dried before treating with 10mg/ml RNase solution (Qiagen, Hilden, Germany). After washing again, the tissues were incubated with 0.3 µg/ml PNA probe TelC-Cy3 (Panagene, Daejeon, Korea) suspended in 70% Formamide 30% water, 10 mmol/L Tris, pH 7.5, heated to 78ºC for 10 min then left overnight at room temperature. The tissues were then washed sequentially with Formamide buffer (70% Formamide, 30% water with 10 mmol/L Tris, pH 7.5.) then PBS containing 0.1% Tween, blocked with 3% BSA (Sigma, St. Louis, MO), 10% donkey serum and incubated overnight at 4ºC with rabbit anti-CC10 antibody for mouse tissues, E-cadherin (Cell signaling 3195S) and pro-SPC (ab3786) for human tissues. Tissues were washed with PBS containing 0.1% Tween and incubated with donkey anti-goat Alexa 488 secondary antibody at 20ºC for 1 h, washed, and mounted using prolong gold anti-fade mounting medium with DAPI (Life Technologies). Images were acquired using a Zeiss Axio Imager 2 microscope (Zeiss, Oberkochen, Germany) and telomere signal intensity quantified using MetaMorph imaging analysis software (Molecular Devices, Sunnyvale, CA). The signal intensity ratio of the Cy3-labeled telomere peptide nucleic acid (PNA) probe and the nuclear stain DAPI correlates to telomere length.

**Quantitative PCR:** Total RNA was extracted from homogenized lung tissues suspended in Qiazol lysis reagent (Qiagen, 79306) according to manufacturer's instructions using miRNeasy Mini Kit (Qiagen, 217004). Isolated RNA was used to synthesize complementary DNA (cDNA) with High-Capacity cDNA Reverse Transcription kit (Applied Biosystems, 4368814) Real time PCR was performed on VIIA-7 (Applied Biosystems). The amplification program was 95ºC for 2 min followed by 40 cycles of combination of 95ºC for 15 sec, 60ºC for 30 sec and 72ºC for 1 min. Relative mRNA levels of target genes were normalized to B-actin mRNA expression and analyzed by $2^{-\Delta\Delta ct}$ method. The sequences of mouse primers are as follows: SMA: forward, 5'-CTCTCTTCCAGCCATCTTTCAT-3', reverse, 5'-TATAGGTGGTTTCGTGGATGC-3'; Collagen1a1 5'-TAGGCCATTGTGTATGCAG -3', reverse, 5'-ACATGTTCAGCTTTGTGGACC-3'; SPC: forward, 5'-ATGGACATGAGTAGCAAAGAGGT-3', reverse, 5'-CACGATGAGAAGGCGTTTGAG -3'; CC10: forward 5'-ATGAAGATCGCCATCACAATCAC-3', reverse 5'-GGATGCCACATAACCAGACTCT-3'; B-actin: forward 5'-CTAAGGCCAACCGTGAAAAG-3', reverse 5'-ACCAGAGGCATACAGGGACA-3'.

**Flow cytometry:** Murine lungs were perfused free of blood, digested in Dispase (50U/mL) for 45 min at 37°C in water bath and suspended in DMEM containing 10 % FBS and DNase I. Cell suspension was filtered through 70 µm and 40 µm filters. Red cell lysis buffer was used to eliminate any RBCs in cell suspension. Cell count was performed. Cells were incubated with a cocktail of conjugated fluorescent antibodies in blocking buffer containing 1% BSA for 30 min at 4°C. After washing, cells were suspended in buffer containing 1% BSA and analyzed on an LSR II flow cytometer. Data were analyzed on FlowJO software V10 (Ashland, OR). The following anti-mouse antibodies were purchased from Biolegend: Percp Cy5.5 CD45 (Catalog no. 103132), PE Cy7 CD3 (Catalog no. 100219), FITC NK1.1 (Catalog no. 108705) and a fixable viability antibody Aqua eFluor780 (ebioscience; catalog no. 65-0865-14).

**Enzyme-Linked Immunosorbent Assay analysis (ELISA):** BAL fluid supernatant was used to determine active TGF-β1 (Biolegend, 437707), IL-6 (eBioscience, 88-7064) and CCL2 Duo Set (R&D systems, DY479-05) ELISA according to the manufacturer's instructions.

**Senescence Associated-β-galactosidase (SA-β-gal) assay:** To test for SA-β-gal (4) staining on tissues following protocol was used as described (5). Briefly, frozen sections were thawed on ice and fixation solution consisting of 2% formaldehyde and 0.2% glutaraldehyde was dispensed. After 2 min of fixation, slides were washed in PBS twice and air-dried. Staining solution was applied on tissue sections and left in a non-Co2 incubator at 37$^{o}$C for 16 hours. To stain senescent cells by cytochemical method 5-bromo-4-chloro-3-indolyl-β-D-galactoside (X-gal) 1 mg/mL was used as a substrate.

**Lung Mechanics:** Lung mechanics were measured as described by the manufacturer (FlexiVent; SCIREQ Inc, Canada). To measure pulmonary physiology mice were anesthetized with i.p. ketamine/xylazine and adequate depth of anesthesia was assured by observing pedal withdrawal reflex. A tracheostomy was performed on the mouse in deep-state anesthesia. A tubing adaptor (20 gauge) was used to cannulate the trachea. The mouse was then attached to a rodent ventilator and pulmonary mechanics analyzer and ventilated at a tidal volume of 9 ml/kg, a frequency of 150 breaths/minute, and 3 cm H2O positive end-expiratory pressure. The mouse was paralyzed with pancuronium bromide (0.1 mg/kg intraperitoneally) to facilitate static compliance measurement. before tracheal cannulation and pancuronium administration. Lung mechanics were measured using forced oscillation maneuver algorithms using flexivent software. Briefly, a deep inflation of subject's lungs to 30 cm H2O was done over a period of 3 seconds to measure inspiratory capacity.

A single frequency forced oscillation maneuver (Snapshot-150 which is a 2.5Hz perturbation to match 150 breath/minute mouse ventilation rate) was used to calculate parameters of entire respiratory system such as Resistance (R), Compliance (C) and Elastance (E). A broadband forced oscillation maneuver was used with a Quicktime-3 perturbation over a wide range of frequencies. Constant phase model (6) is a mathematical model employed to calculate Newtonian resistance ($R_N$), tissue damping (G) and tissue elastance (H) parameters. Pressure-Volume curves utilize stepwise changes in pressure or volume. Values are recorded at the end of each step. From the PV-loop values, Salazar-Knowles equation derives static compliance (Cst), Parameter A (an estimate of inspiratory capacity) and shape constant (K).

**MicroCT imaging:** Mouse lungs were perfused free of blood and fixed in 4 % PFA at $4^oC$ overnight. Lungs were washed in PBS 4 times 30 min each. Lungs were dehydrated in 25 % Lugol's staining solution at $4^oC$ for 48 h. Lungs were embedded in 3 % agarose. Lung specimens were scanned on a Scanco Medical µCT 50 high-resolution specimen scanner immediately after being embedded in agarose gel. The energy of the X-ray was set at 45 kVp and 88 µA. Specimens were scanned at a resolution of 10-µm voxel size and using an integration time of 500 milliseconds. Quantitative analyses were performed using Scanco µCT Evaluation Program v6.0. The region of interest was manually delineated at each slice to include the entire lung area. The delineated slices were combined to create a volume, which was then analyzed for three-dimensional volumetric and morphometric indices. An upper threshold of 160 grayscale units, out of 1000, was applied to segregate lung airspace and lung tissue. A Gaussian filter with sigma of 0.8 and support of 1 was applied to remove background noise. 3D-rendered segmented images were generated using Scanco Ray v4.0.

**Statistics**: Statistical analysis were performed with GraphPad Prism software (Version 7.0) P values were calculated using two-tailed Student's *t* test. Welch's correction was applied for analysis of unpaired groups. Differences in survival were analyzed by *log-rank (Mantel-Cox)* test. Data are represented as the mean ± SEM. *P* values represent the significance using the following symbols. ns = not significant, p>0.05, *p<0.05, **p<0.01, ***p<0.001, ****p<0.0001.
